# Untargeted cord blood metabolomics reveals altered lipid metabolism in neonates with gastroschisis

**DOI:** 10.64898/2026.06.04.730243

**Authors:** Hsuan-Yuan Wang, Bryan T. Oshiro, Neela Rahseparian, Lauren Crabtree, Joshua F. Robinson, Stephanie L Gaw, Ciprian Gheorghe

**Author notes:** Co-corresponding authors: Stephanie L. Gaw, MD, PhD, Associate Professor, Division of Maternal-Fetal Medicine Department of Obstetrics, Gynecology and Reproductive Sciences University of California, San Francisco, 513 Parnassus Ave, Box 0556 16HSE San Francisco, CA 94143, Ciprian Gheorghe, MD, PhD, Division of Maternal-Fetal Medicine, Department of Gynecology and Obstetrics Loma Linda University School of Medicine, 11234 Anderson Street, Loma Linda, CA 92354.

## Abstract

Gastroschisis is a congenital abdominal wall defect in which fetal intestines herniate into the amniotic cavity. Despite 97% surgical repair success rate, 40% of affected infants require hospital readmission due to gastrointestinal complications, where underlying mechanisms remain poorly characterized. We hypopthesized that the cord blood metabolome of neonates with gastroschisis differs systematically from controls and may reveal pathway-level alterations relevant to neonatal physiology. Cord blood plasma collected at delivery (23 samples each group) was analyzed using ultra-performance liquid chromatography coupled with tandem mass spectrometry. Unsupervised principal component analysis and hierarchical clustering demonstrated significant separation between groups (PERMANOVA *pseudo-F* = 4.632, *R²* = 0.095, *p* = 0.001). 53 metabolites met criteria for differential abundance, 75% were lipids. Key alterations included reduced free fatty acids, increased fatty acid amides and ceramides, disrupted steroid and bile acid metabolism, and decreased biliverdin and bilirubin isomers. Our findings provide insight into gastroschisis pathophysiology and identify potential biomarkers for future investigation.

## Introduction

Gastroschisis is a congenital anomaly characterized by a full thickness defect in the abdominal wall, typically to the right of the umbilical cord insertion, through which the fetal intestines herniate into the amniotic cavity. The global prevalence has been estimated at 1.79 per 10,000 births, with notable geographic variation; North and South America exhibit the highest rates, and Europe and Asia the lowest^1,2^. In the United States, the prevalence was estimated at 4.3 per 10,000 live births during 2012 to 2016^3^, with a subsequent decline observed from 2016 to 2022^4^. Identified risk factors include young maternal age^5^, tobacco or alcohol exposure during early pregnancy^6^, genitourinary infection in early gestation^7^, familial recurrence^8,9^, and prenatal cannabis exposure^10,11^.

Although postnatal survival in high resource settings exceeds 97%^12,13^, hospital readmission affects up to 40% of affected infants and is driven primarily by bowel obstruction, abdominal distention, and gastroenteritis^14,15^. Prenatal detection rates exceed 80% on second trimester ultrasonography in high resource settings^16,17^, and standard management consists of urgent neonatal surgical closure with subsequent staged reduction of viscera to the abdominal cavity^18,19^. Despite the high success rate of surgical closure, the metabolic and physiological derangements that predispose to long term complications in this population remain incompletely understood.

Cord blood plasma sampled at delivery contains a mixture of metabolites of fetal origin, derived from intrinsic fetal metabolism, and metabolites of maternal origin, transferred across the placenta^20^. Cord blood metabolomic profiling has proven informative in characterizing congenital conditions and fetal growth restriction^21,22^, environmental exposures^23^, and developmental physiology^24^. The technique therefore offers a tractable approach to interrogate the metabolic milieu of the gastroschisis newborn at delivery.

We thus tested the hypothesis that the cord blood metabolomic profile of neonates with gastroschisis differs systematically from that of healthy controls, and that these differences may identify pathway-level alterations relevant to the clinical phenotype. To test this hypothesis, we performed untargeted ultra performance liquid chromatography tandem mass spectrometry on cord blood plasma collected at delivery from 23 neonates with gastroschisis and 23 healthy controls.

## Results

### Demographic and clinical characteristics

The clinical characteristics of the study participants are summarized in Table 1. Maternal age was lower in the gastroschisis group (23.3 ± 2.4 vs. 29.3 ± 5.5 years, p < 0.0001), consistent with the previously reported association between young maternal age and gastroschisis^25,26^. None of the gastroschisis cases occurred in mothers of advanced maternal age, compared with 2 (8.7%) of controls (*p* = 0.489). Body mass index did not differ between groups (30.0 ± 6.4 vs. 28.9 ± 4.7 kg/m², *p* = 0.539), and the prevalence of obesity was similar (30.4% vs. 39.1%, *p* = 0.758). Racial and ethnic composition was comparable between groups (*p* = 0.458), with the majority of participants identifying as Hispanic (69.6% gastroschisis vs. 56.5% control). Gestational age at delivery was lower in the gastroschisis group (36.6 ± 1.9 vs. 38.4 ± 2.5 weeks, p = 0.009), although the rate of preterm birth did not differ (26.1% vs. 17.4%, *p* = 0.722). Maternal comorbidities were uncommon and balanced across groups. Chronic hypertension, pregestational diabetes, and gestational diabetes were absent from both cohorts, and maternal smoking, sexually transmitted infection (chlamydia or gonorrhea) in pregnancy, preterm labor, preterm premature rupture of membranes, and chorioamnionitis occurred at low and statistically indistinguishable rates (Table 1). A single control pregnancy was complicated by gestational hypertension, with no analog among cases. Primigravidity was similar between groups (56.5% vs. 39.1%, *p* = 0.376), as was the distribution of delivery mode (*p* = 0.734) and the proportion of male infants (47.8% in both groups). Neonatal characteristics, in contrast, differed substantially. Birthweight was lower in the gastroschisis group (2503 ± 515 vs. 3032 ± 585 g, *p* = 0.002), and the proportion classified as small for gestational age was nearly fourfold higher (52.2% vs. 13.0%, *p* = 0.011), consistent with the established association between gastroschisis and fetal growth restriction. Cases demonstrated lower 1-minute and 5-minute Apgar scores than controls (median 7 versus 8 at 1 minute, *p* = 0.002; median 9 versus 9 at 5 minutes, *p* = 0.033), and 15 of 23 cases (65 %) required endotracheal intubation during the neonatal admission compared with none of 23 controls. Neonatal intensive care unit length of stay was markedly longer in cases (median 35 vs. 2 days, <u>p</u> < 0.0001), reflecting the routine need for staged abdominal wall closure and prolonged parenteral nutrition. Severe neonatal complications, defined as necrotizing enterocolitis, treated clinical or culture positive sepsis, bowel resection, packed red blood cell transfusion, high frequency oscillatory ventilation, treated pulmonary hypertension, or vasopressor support, occurred in 11 of 23 gastroschisis cases (47.8%) compared with 1 of 23 controls (4.3%, *p* = 0.002).

**Table 1.**
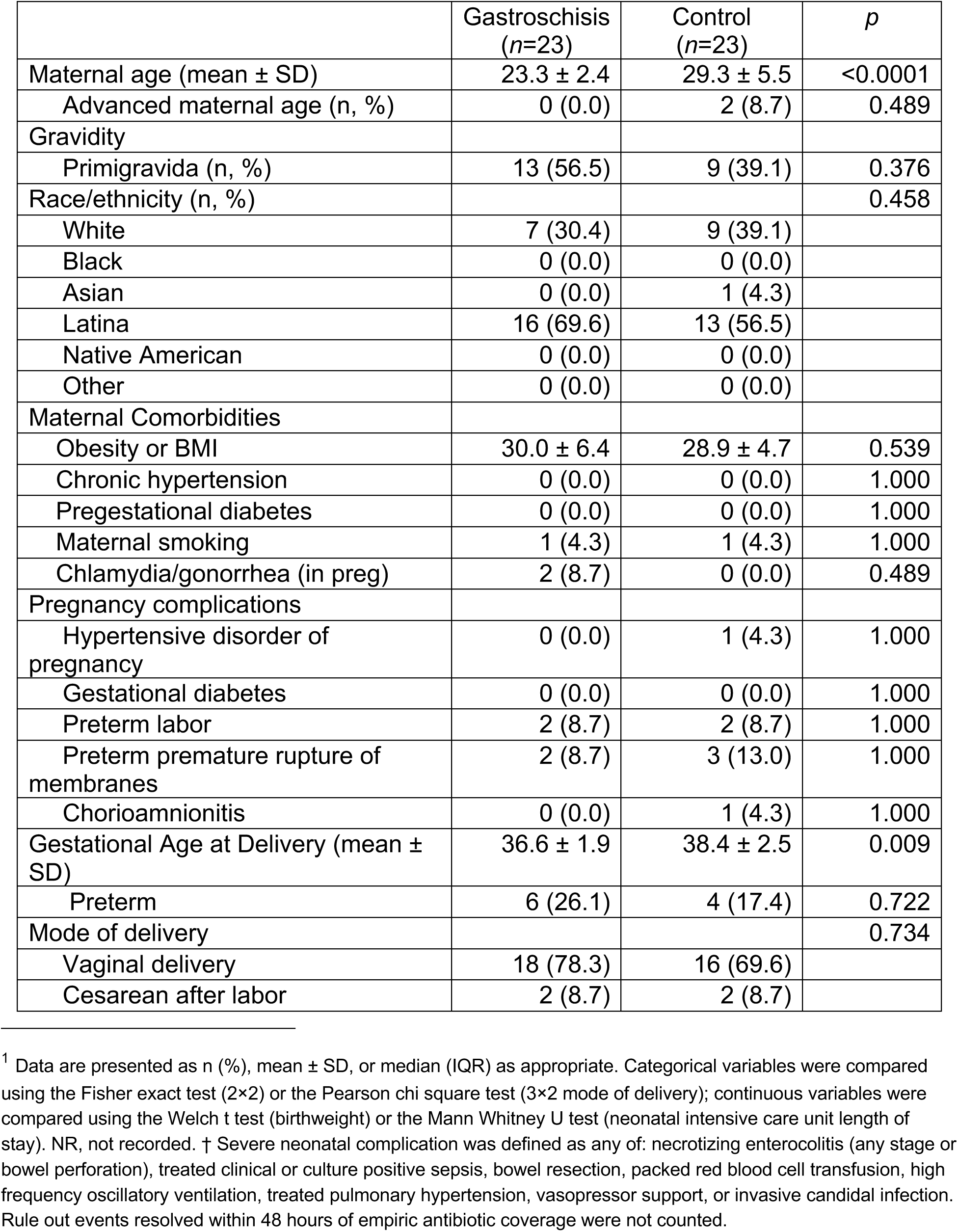
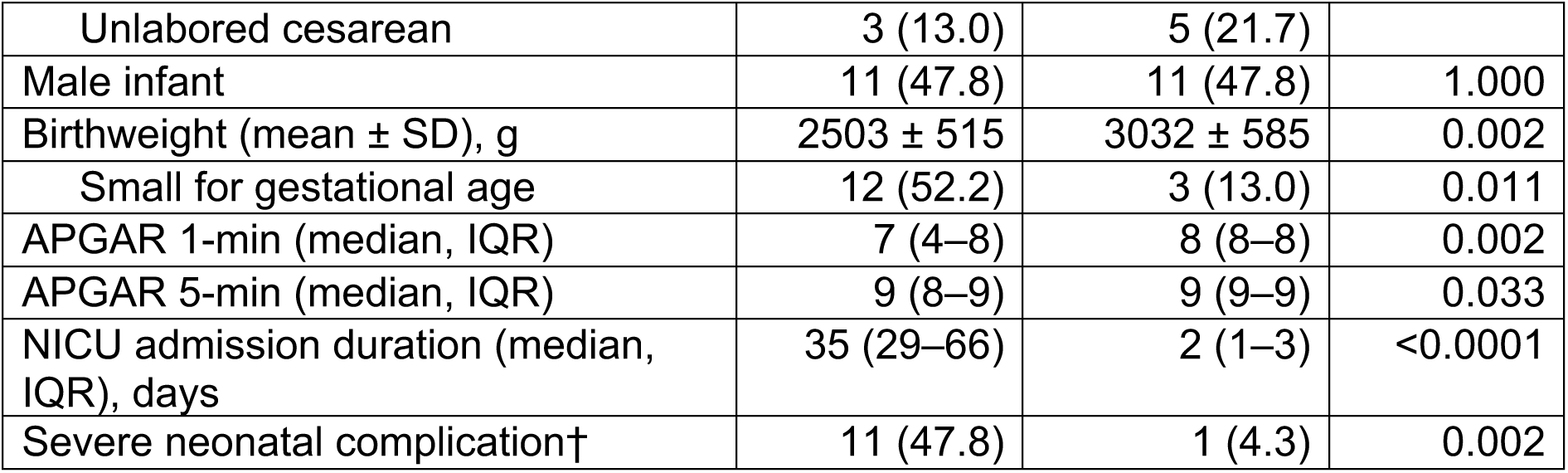
Participant characteristics and perinatal outcomes^1^.

|  | Gastroschisis<br>(n=23) | Control<br>(n=23) | *p* |
| --- | --- | --- | --- |
| Maternal age (mean ± SD) | 23.3 ± 2.4 | 29.3 ± 5.5 | <0.0001 |
| Advanced maternal age (n, %) | 0 (0.0) | 2 (8.7) | 0.489 |
| Gravidity |  |  |  |
| Primigravida (n, %) | 13 (56.5) | 9 (39.1) | 0.376 |
| Race/ethnicity (n, %) |  |  | 0.458 |
| White | 7 (30.4) | 9 (39.1) |  |
| Black | 0 (0.0) | 0 (0.0) |  |
| Asian | 0 (0.0) | 1 (4.3) |  |
| Latina | 16 (69.6) | 13 (56.5) |  |
| Native American | 0 (0.0) | 0 (0.0) |  |
| Other | 0 (0.0) | 0 (0.0) |  |
| Maternal Comorbidities |  |  |  |
| Obesity or BMI | 30.0 ± 6.4 | 28.9 ± 4.7 | 0.539 |
| Chronic hypertension | 0 (0.0) | 0 (0.0) | 1.000 |
| Pregestational diabetes | 0 (0.0) | 0 (0.0) | 1.000 |
| Maternal smoking | 1 (4.3) | 1 (4.3) | 1.000 |
| Chlamydia/gonorrhea (in preg) | 2 (8.7) | 0 (0.0) | 0.489 |
| Pregnancy complications |  |  |  |
| Hypertensive disorder of pregnancy | 0 (0.0) | 1 (4.3) | 1.000 |
| Gestational diabetes | 0 (0.0) | 0 (0.0) | 1.000 |
| Preterm labor | 2 (8.7) | 2 (8.7) | 1.000 |
| Preterm premature rupture of membranes | 2 (8.7) | 3 (13.0) | 1.000 |
| Chorioamnionitis | 0 (0.0) | 1 (4.3) | 1.000 |
| Gestational Age at Delivery (mean ± SD) | 36.6 ± 1.9 | 38.4 ± 2.5 | 0.009 |
| Preterm | 6 (26.1) | 4 (17.4) | 0.722 |
| Mode of delivery |  |  | 0.734 |
| Vaginal delivery | 18 (78.3) | 16 (69.6) |  |
| Cesarean after labor | 2 (8.7) | 2 (8.7) |  |
| Unlabored cesarean | 3 (13.0) | 5 (21.7) |  |
| Male infant | 11 (47.8) | 11 (47.8) | 1.000 |
| Birthweight (mean ± SD), g | 2503 ± 515 | 3032 ± 585 | 0.002 |
| Small for gestational age | 12 (52.2) | 3 (13.0) | 0.011 |
| APGAR 1-min (median, IQR) | 7 (4–8) | 8 (8–8) | 0.002 |
| APGAR 5-min (median, IQR) | 9 (8–9) | 9 (9–9) | 0.033 |
| NICU admission duration (median, IQR), days | 35 (29–66) | 2 (1–3) | <0.0001 |
| Severe neonatal complication† | 11 (47.8) | 1 (4.3) | 0.002 |

### Unsupervised metabolomic profiling

Cord blood plasma underwent untargeted UPLC-MS/MS metabolomic profiling (Fig. 1A). A total of 925 metabolites with confirmed identities were detected. To focus on endogenous biology, 108 xenobiotic compounds were excluded (Supplementary Fig. 1), and metabolites detected in fewer than 70% of the cohort were further excluded, yielding 767 metabolites for downstream analysis. Lipids constituted 57.5% and amino acids 26.1% of the retained metabolites (Fig. 1B). Unsupervised principal component analysis demonstrated partial separation between groups (Fig. 1C), and PERMANOVA confirmed a significant difference in distribution (*pseudo-F* = 4.632, *R²* = 0.095, *p* = 0.001). Hierarchical clustering identified two principal clusters consistent with case status (Fig. 1D), indicating that the cord blood metabolomic profile of neonates with gastroschisis is distinguishable from that of healthy controls.

**Figure 1.**
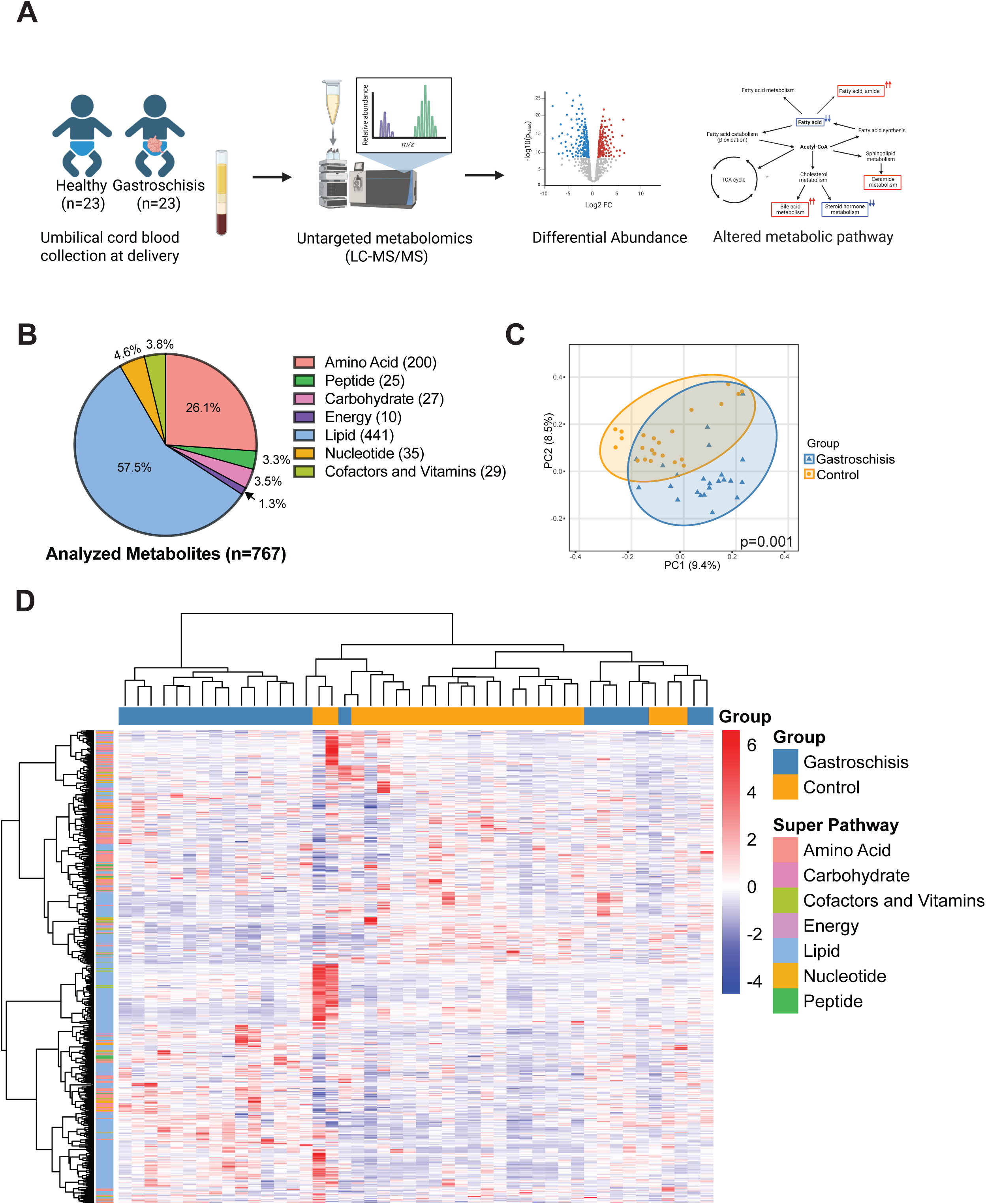
Cord blood metabolomic profiling of gastroschisis and healthy individuals. (A) Study design. Cord plasma samples from neonates with gastroschisis (n = 23) and healthy controls (n = 23) were analyzed using untargeted metabolomics via liquid chromatography tandem mass spectrometry (LC MS/MS). Differential metabolite abundance was assessed, and significantly altered metabolic pathways between the two groups were identified. (B) Categories of the final analyzed metabolites (n = 767). (C) Unsupervised PCA on total analyzed metabolites between gastroschisis and control groups; PERMANOVA pseudo-F = 4.632, R² = 0.09535, p = 0.001. Steel blue triangle: gastroschisis; orange circle: control. (D) Unsupervised heatmap clustering. Metabolites are categorized by superpathway (left Y axis); each column represents one individual from the gastroschisis (steel blue) or control (orange) group. Red indicates higher than average expression and blue indicates lower than average expression.

### Key metabolic pathway alterations

Of the 767 metabolites analyzed, 53 metabolites of confirmed endogenous identity met the differential abundance criteria of fold change (FC) greater than 2 or less than 0.5 at p < 0.05 (Fig. 2A). These 53 metabolites constituted the core metabolic signature examined in subsequent analyses. Lipids were the predominant contributor (75.5%), followed by amino acids (13.2%), cofactors and vitamins (7.5%), and nucleotides and peptides (1.9% each) (Fig. 2B). Mapping of these metabolites to their respective sub pathways revealed that significant changes were uniformly directional within most sub pathways, indicating pathway level rather than metabolite level alterations (Fig. 2C). Sub pathways with greater than 30% activation included fatty acid amide metabolism, dihydroceramide metabolism, and secondary bile acid metabolism, whereas androgenic and estrogenic steroid pathways and hemoglobin and porphyrin metabolism each demonstrated greater than 30% deactivation.

**Figure 2.**
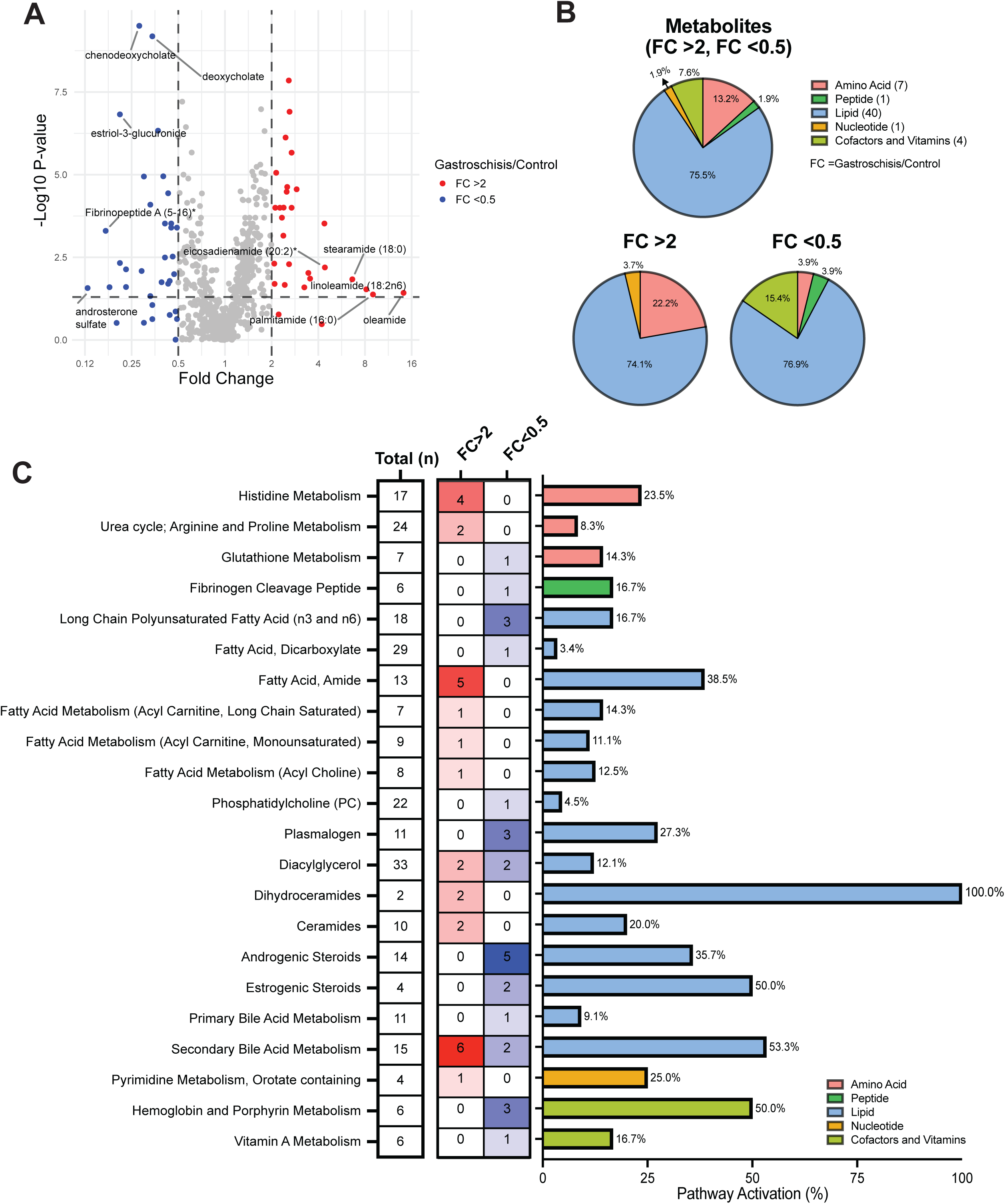
Key metabolites distinguishing neonates with gastroschisis and healthy controls. (A) Volcano plot of all analyzed metabolites (n = 767), with x axis representing log₂ fold change and y axis representing −log₁₀(p value). Metabolites with fold change >2 are highlighted in red and <0.5 in blue (p < 0.05). A total of 53 metabolites met these criteria. (B) Pathway category distribution of the 53 significant metabolites: amino acids (red), peptides (green), lipids (blue), nucleotides (orange), cofactors and vitamins (light green). (C) Pathway activation. Metabolites with differential abundance >2 fold or <0.5 fold were categorized by subpathway. Total (n) is the number of metabolites assigned to each pathway in the reference library. FC > 2 column represents upregulated metabolites; FC < 0.5 column represents downregulated metabolites. Pathway activation (%) is the proportion of significant metabolites relative to the total.

### Sensitivity of the metabolite signature to gestational age

Gastroschisis is associated with both preterm delivery and fetal growth restriction, and our cohort reflected this clinical reality. Cases were delivered earlier than controls (36.6 versus 38.4 weeks) and were more frequently small for gestational age (52.2% versus 13.0%; Table 1). To test whether the cohort metabolite signature was an artifact of prematurity, we repeated the case versus control comparison after restricting to term births, excluding the 6 preterm cases and 4 preterm controls and recomputing the fold change and significance in the resulting 17 cases and 19 controls. The fold change estimator and the significance criterion were identical to the discovery analysis, namely a mean ratio above 2 or below 0.5 together with a Welch t test p value below 0.05, so retention in the term subset is directly comparable to the original selection.

Restriction to term births does not equalize gestational age, because gastroschisis is delivered near 37 weeks under institutional protocol while term controls reach 39 to 40 weeks; within the term subset the cases remained earlier than the controls (37.5 versus 39.3 weeks) and the imbalance in small for gestational age status widened (47% versus 5%). With that caveat, 35 of the 53 cohort significant metabolites remained significant when the comparison was restricted to term births (Fig. 3; Supplementary Table 1), indicating that most of the signature is not an artifact of prematurity. The retained metabolites included all the heme and bilirubin species, the ceramides, the primary bile acid, and the majority of the secondary bile acids and androgenic steroids. The 18 metabolites that lost significance fell into two groups. For six, the mean fold change itself attenuated toward the null in term births, consistent with a contribution from prematurity or growth restriction, and this group included the urea cycle metabolites. For the remaining twelve, including the fatty acid amides that showed the largest unadjusted elevations, oleamide, palmitamide, linoleamide, and stearamide, the mean fold change remained large but no longer reached significance in the smaller subset, with only eicosadienamide among the fatty acid amides retained. The fatty acid amide elevations in particular are driven by a few infants rather than by a uniform shift, with a median case to control ratio near one, and are not explained by gestational age, because the mean ratio was essentially unchanged when preterm infants were excluded. The cohort level fold changes of the fatty acid amides should therefore be interpreted with caution.

**Figure 3.**
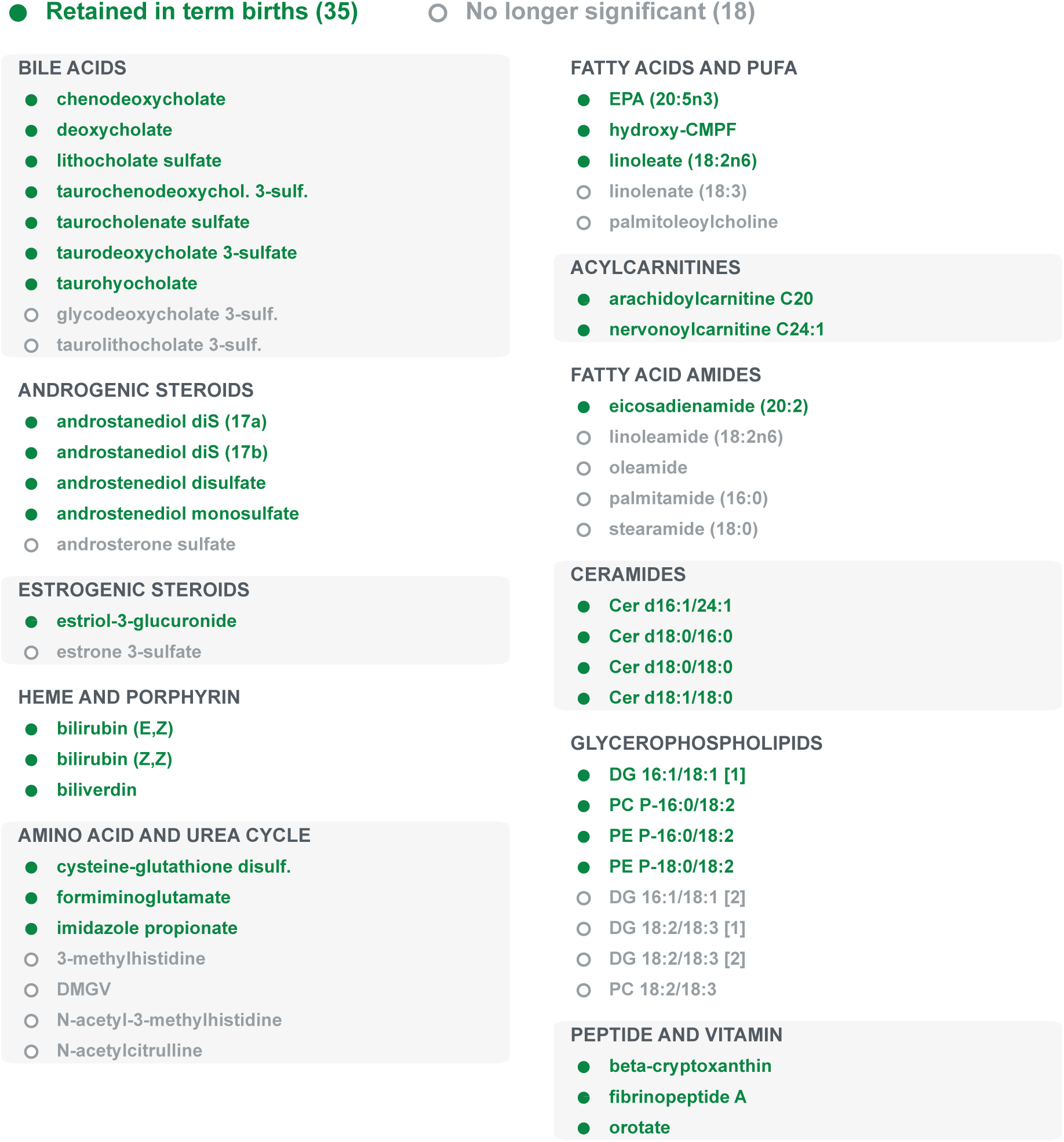
Robustness of the gastroschisis metabolite signature to gestational age. Each of the 53 cohort significant metabolites is shown grouped by biological pathway and colored by whether it remained significant when the case versus control comparison was restricted to term births (17 cases versus 19 controls), using the same criterion that defined the 53 (FC > 2 or < 0.5 with a p value below 0.05). Filled green markers denote the 35 metabolites retained in term births; open grey markers denote the 18 that were no longer significant. The fatty acid amides, whose unadjusted elevations were the largest in the cohort, were largely not retained; their mean fold change persisted in term births but lost significance, reflecting an elevation driven by a few infants, with a median case to control ratio near one, rather than a gestational age effect.

### Altered lipid metabolism

Because lipids constituted the predominant class of altered metabolites, we performed a global assessment of lipid metabolic pathway alterations centered on the principal intermediates, fatty acid and acetyl-CoA. Cord blood from neonates with gastroschisis exhibited reduced abundance of long chain fatty acids accompanied by increased abundance of metabolites in downstream pathways, including fatty acid metabolism and fatty acid amide metabolism (Fig. 3A; Supplementary Table 2 and 3). Within phospholipid metabolism (Supplementary Table 4), gastroschisis samples showed reduced plasmalogen levels, a reduction in a polyunsaturated phosphatidylcholine species (1-linoleoyl-2-linolenoyl-GPC, FC = 0.23), and mixed alterations in diacylglycerol centered pathways characterized by reduced polyunsaturated and increased monounsaturated species. Acetyl-CoA derived sphingolipid metabolism was activated, with elevated dihydroceramides and ceramides. In cholesterol derived pathways, steroid hormone metabolism was attenuated whereas bile acid metabolism was augmented. Overall, gastroschisis was associated with global lipid pathway remodeling characterized by elevated fatty acid amides, sphingolipids, ceramides, bile acids, and reduced fatty acids and steroid hormones (Fig. 3B).

### Increased fatty acid amide and ceramide metabolism

Fatty acid amides are bioactive lipids generated from free fatty acids by N-acyltransferases and degraded by fatty acid amide hydrolase (FAAH)^27^ (Fig. 4A). All five fatty acid amides meeting the prespecified detection threshold were elevated in the gastroschisis group (Fig. 4B and C), with oleamide demonstrating a 14-fold increase, palmitamide a nine-fold increase, linoleamide an eight-fold increase, stearamide a seven-fold increase, and eicosadienamide a four-fold increase. The magnitude of these elevations is consistent with reduced FAAH mediated hydrolysis or enhanced N-acyltransferase activity. Ceramide biosynthesis proceeds through three principal routes: *de novo* synthesis from palmitoyl-CoA and serine, sphingomyelin hydrolysis, and the sphingosine salvage pathway (Fig. 4D). In our cohort, both dihydroceramides and ceramides were elevated (Fig. 4E and F; Supplementary Table 5), consistent with enhanced *de novo* sphingolipid biosynthesis or impaired catabolism within the sphingomyelin or salvage pathways.

**Figure 4.**
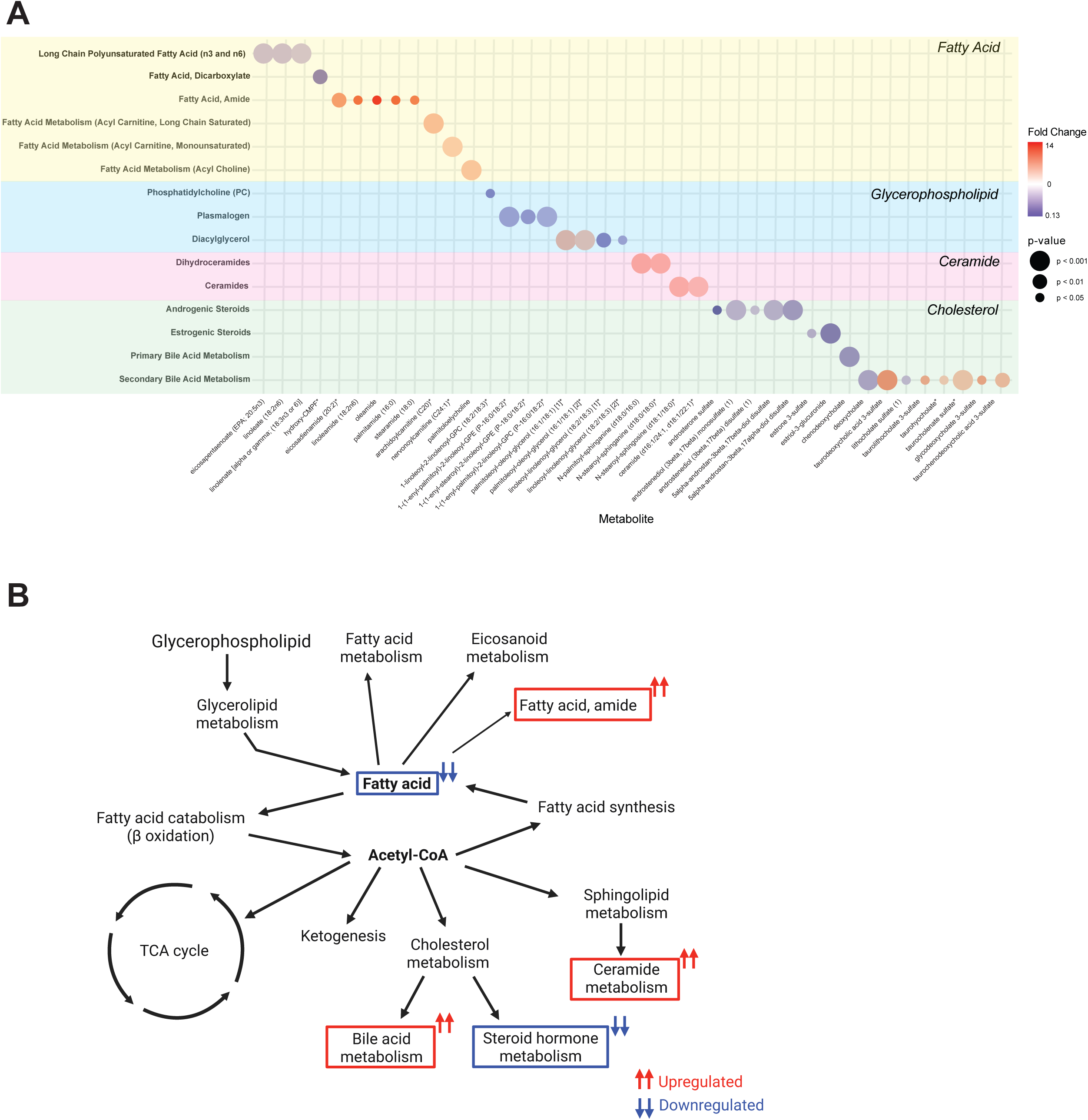
Lipid metabolite changes in neonates with gastroschisis. (A) Significant lipid metabolites (p < 0.05; FC >2 or <0.5) by lipid subpathway: fatty acids (yellow), glycerophospholipids (blue), ceramides (pink), and cholesterol (green). Bubble color reflects fold change (red, increased; blue, decreased) and bubble size reflects significance (p value). (B) Schematic of lipid metabolism with key altered pathways highlighted. Red indicates overall upregulation and blue indicates overall downregulation in gastroschisis.

### Decreased steroid hormone and increased bile acid metabolism

Steroid hormone and bile acid metabolism share cholesterol as their common precursor. Steroid hormone biosynthesis comprises androgenic and estrogenic arms (Fig. 5A), beginning with conversion of cholesterol to dehydroepiandrosterone (DHEA) via pregnenolone. DHEA serves as the precursor for several androgenic intermediates and end products including DHEA sulfate (DHEA-S), 16α hydroxy DHEAS, androstanediol, androsterone, and testosterone. Androstenedione and testosterone may serve as substrates for estrogen biosynthesis, yielding estrone sulfate, estriol sulfates, and glucuronide conjugates. Both androgenic and estrogenic steroids were globally reduced in neonates with gastroschisis (Fig. 5B and C; Supplementary Table 6), suggesting a shift in cholesterol utilization toward alternative pathways.

**Figure 5.**
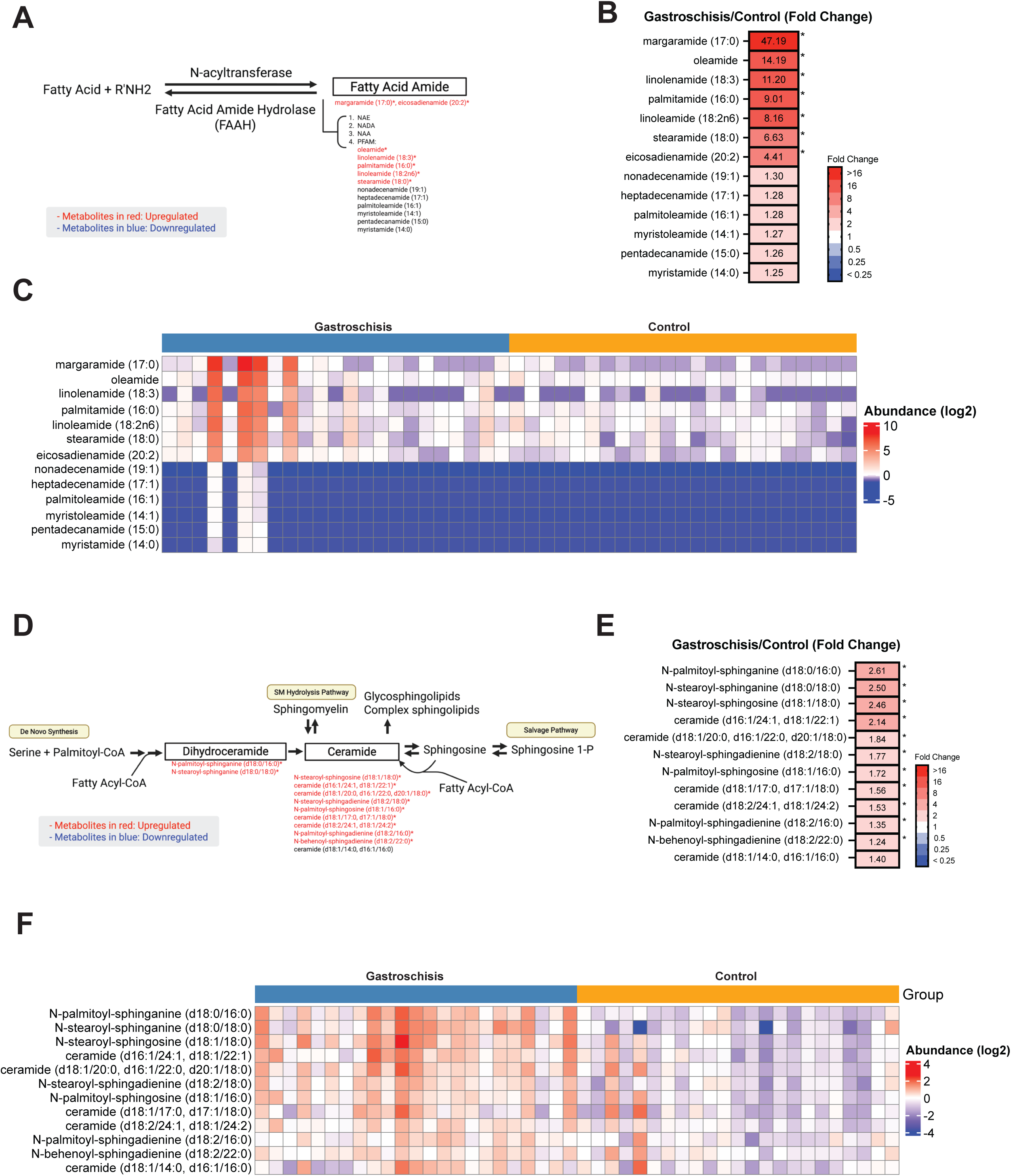
Highly upregulated fatty acid amides and ceramides in neonates with gastroschisis. (A) Schematic of fatty acid amide metabolism showing synthesis via N-acyltransferase and degradation by FAAH. Red, upregulated; blue, downregulated; asterisks denote p < 0.05. (B) Fold change of identified fatty acid amides ranked by magnitude. (C) Heatmap of log₂ transformed abundance per sample, grouped by condition. (D) Schematic of ceramide metabolism with identified metabolites annotated. (E) Fold change of ceramide metabolites ranked by magnitude. (F) Heatmap of log₂ transformed abundance per sample, grouped by condition.

Bile acid metabolism comprises hepatic primary and intestinal secondary stages^28^ (Fig. 5D). In the liver, cholesterol is converted to the primary bile acids cholic acid and chenodeoxycholic acid, which are conjugated with taurine or glycine to form taurocholate, glycocholate, taurochenodeoxycholate, and glycochenodeoxycholate. These conjugates are secreted into the small intestine where gut microbiota generate secondary bile acids and their conjugates. Approximately 95% of bile acids are reabsorbed via enterohepatic circulation, with the remainder excreted in feces^29^. Compared with healthy controls, neonates with gastroschisis exhibited reduced cholic acid and chenodeoxycholic acid but elevated taurocholate, indicating activation of primary bile acid metabolism (Fig. 5E and F; Supplementary Table 7). The majority of glucuronidated and sulfated bile acids were elevated approximately two to four-fold, with the exception of lithocholate sulfate, which was reduced. The accumulation of conjugated and sulfated species in the circulation is consistent with compromised enterohepatic circulation.

### Reduced hemoglobin and bilirubin metabolism

Heme is produced predominantly in the liver and bone marrow, with key biosynthetic steps localized to the mitochondria^30^ (Fig. 6A). Heme oxygenase converts heme to biliverdin^31^, which is reduced to bilirubin by biliverdin reductase^32^. Bilirubin exists in four structural isomers (ZZ, ZE, EZ, EE)^33^. Unconjugated bilirubin binds albumin in the plasma and is transported to the liver, where it is conjugated with glucuronic acid for biliary excretion^34^. Within the intestine, gut microbiota converts conjugated bilirubin to urobilinogen, which is either further metabolized to stercobilin and excreted in feces or reabsorbed and excreted as urobilin in urine^35^. In our cohort, biliverdin and all bilirubin isomers were reduced, with a non-significant trend toward heme accumulation (Fig. 6B and C), consistent with attenuated heme oxygenase activity.

**Figure 6.**
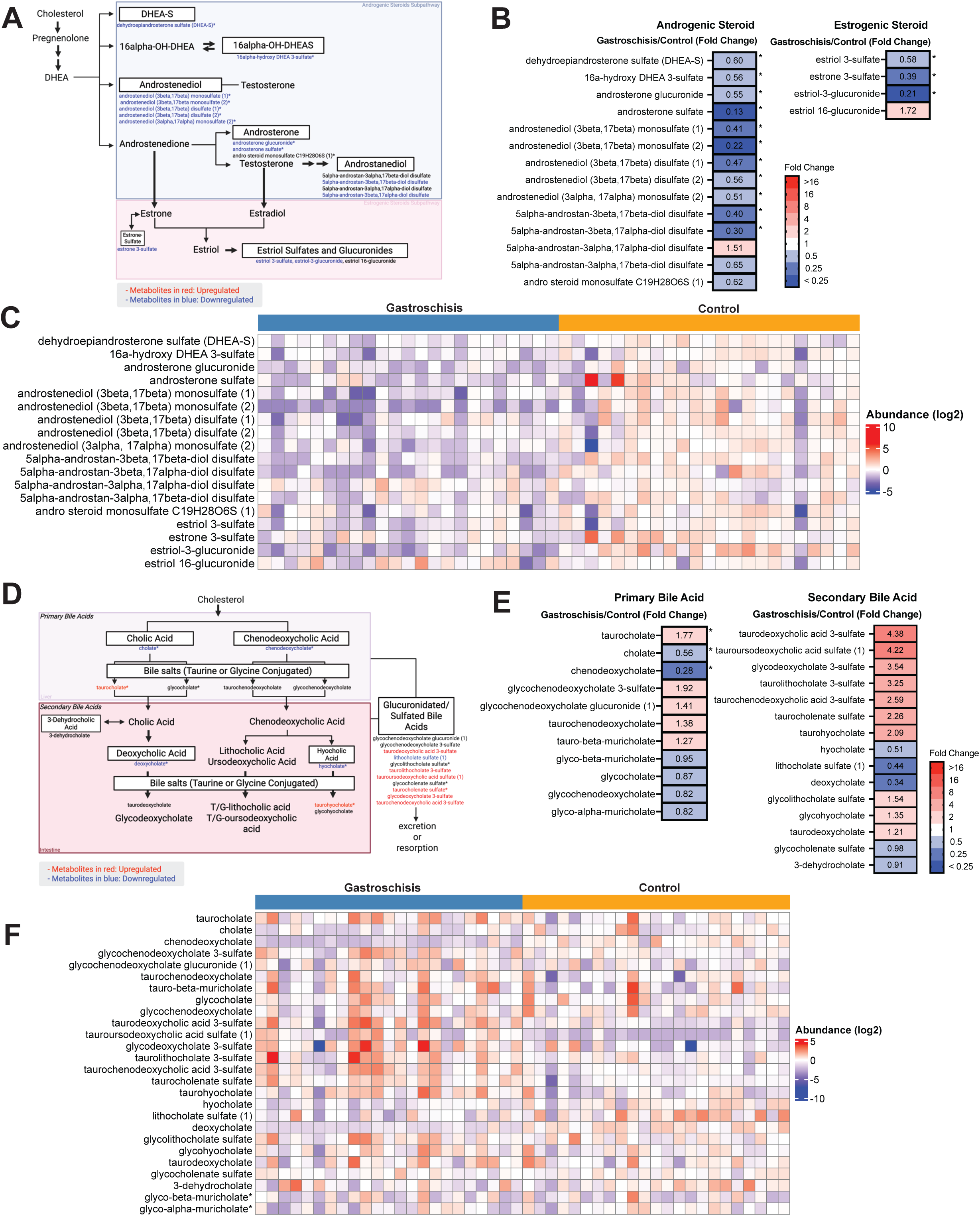
Reduced steroid hormone metabolism and increased bile acid metabolism in neonates with gastroschisis. (A) Schematic of steroid hormone metabolism with annotated identified metabolites. (B) Fold change of identified steroid hormone metabolites. (C) Heatmap of log₂ transformed abundance per sample. (D) Schematic of bile acid metabolism with annotated metabolites. (E) Fold change of identified bile acid metabolites. (F) Heatmap of log₂ transformed abundance per sample.

## Discussion

The present study constitutes, to our knowledge, the first untargeted metabolomic profiling of cord blood plasma in neonates with gastroschisis. Of the 767 endogenous metabolites detected, 53 displayed differential abundance between cases and controls, and approximately 75% of these were lipids. The principal alterations included a global reduction in free fatty acids, marked elevations in fatty acid amides and ceramides, attenuated steroid hormone biosynthesis, a shift from unconjugated to conjugated bile acids, and reduced biliverdin and bilirubin. These observations are consistent with a previous report describing altered fatty acid composition in maternal and fetal blood from gastroschisis cases^36^ and extend that work to a comprehensive survey of the cord blood metabolome.

Long chain polyunsaturated fatty acids (LC-PUFA) were globally reduced in the gastroschisis group. Because the fetus has limited capacity for endogenous LC-PUFA synthesis, these molecules are predominantly maternal in origin and are transferred across the placenta during the third trimester^37^. Reduced LC-PUFA availability during this interval has been associated with impaired brain and central nervous system development^37^, and neonates with gastroschisis, particularly those with severe phenotype, demonstrate higher rates of neurodevelopmental delay^38^. The reduction in cord blood LC-PUFA observed here is concordant with both the established maternal to fetal transfer literature and the clinical risk profile of these neonates, although the partition between gastroschisis specific effects and effects mediated through earlier delivery and growth restriction cannot be resolved at the present cohort size. Maternal LC-PUFA supplementation during the third trimester may therefore merit prospective evaluation as a prophylactic intervention in pregnancies complicated by gastroschisis, with the recognition that the same intervention may benefit any pregnancy at risk for preterm or growth restricted delivery and is not specific to gastroschisis in its rationale.

Fatty acid amides function as endocannabinoid ligands and signaling molecules in the central nervous system, with roles in sleep regulation, locomotion, and angiogenesis^39^. Their hydrolysis is mediated by fatty acid amide hydrolase (FAAH), which is enriched in metabolically active tissues including brain, liver, pancreas, intestine, and testis^40^. Exogenous cannabinoids compete with anandamide for cannabinoid receptor binding and thereby suppress FAAH activity, leading to fatty acid amide accumulation^41^. Of critical importance, prenatal cannabis exposure has been consistently identified as a risk factor for gastroschisis in epidemiologic studies from California, Canada, and Europe^10,11,42,43^. The elevation of fatty acid amides in cord blood, led by a 14-fold increase in oleamide and shared across all five members of the class meeting our detection threshold (Fig. 4B), may therefore reflect, at least in part, prenatal cannabis exposure or a related disturbance of endocannabinoid signaling. The concurrent reduction in free fatty acid precursors further supports this interpretation. This interpretation should nonetheless be made cautiously, because the fatty acid amide elevations are driven by a small number of infants rather than by a uniform increase, with a median case to control ratio near one. The mean fold change remained large when the comparison was restricted to term births, and was therefore not explained by gestational age, but it no longer reached statistical significance in the smaller subset, with only eicosadienamide retained (Figure 7). Prenatal cannabis exposure was not ascertained in this cohort, and whether the elevation in these few infants reflects maternal cannabis use or another disturbance of endocannabinoid signaling will require a larger cohort with documented exposure in which the distribution of cord blood fatty acid amides can be examined directly.

**Figure 7.**
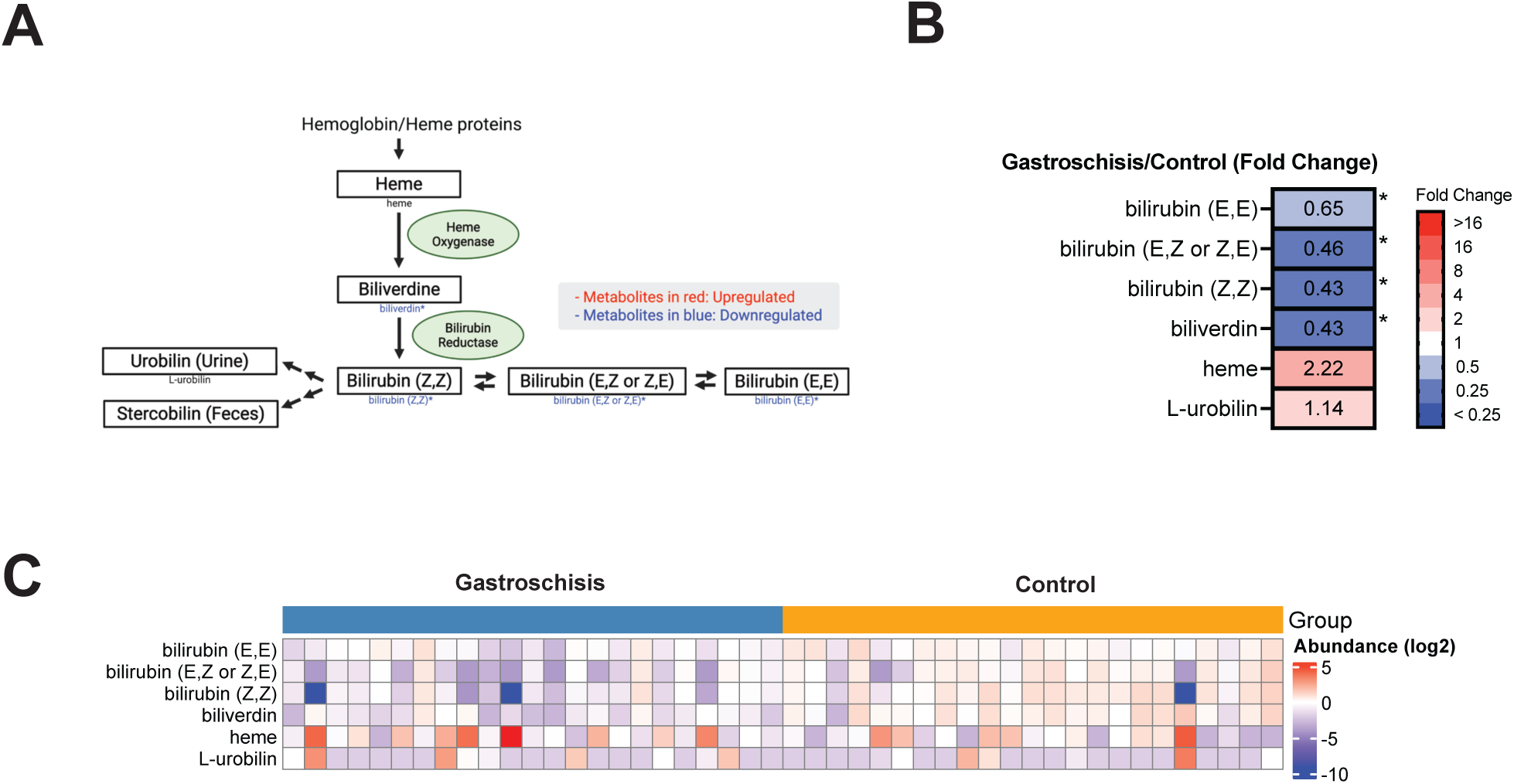
Reduced heme and bilirubin metabolism in neonates with gastroschisis. (A) Schematic of heme and bilirubin metabolism with annotated identified metabolites. (B) Fold change of identified heme and bilirubin metabolites. (C) Heatmap of log₂ transformed abundance per sample, grouped by condition.

Both androgenic and estrogenic steroid pathways were attenuated in the gastroschisis group. Dehydroepiandrosterone sulfate (DHEA-S), produced primarily by the fetal adrenal gland, serves as the principal substrate for placental estrogen synthesis^44,45^. The coordinated reduction of DHEA-S, downstream androgens including testosterone, and estrogen metabolites suggests impaired function of the fetal adrenal placental axis. Of note, approximately one third of male newborns with gastroschisis present with unilateral cryptorchidism, a condition associated with low testosterone^46^, which is congruent with the steroidogenic deficit observed at the metabolomic level. Because steroid maturation in cord blood is itself dependent on gestational age, and the gastroschisis and control groups differed in this dimension, the relative contributions of a gastroschisis specific adrenal to placental axis disturbance and of gestational age dependent fetal adrenal maturation cannot be disentangled at the present cohort size^47^.

Bile acid metabolism is initiated in the fetal liver, with conjugated species predominantly of fetal origin and glycocholate of maternal origin^48,49^. Neonates with gastroschisis exhibited a near twofold elevation of taurocholate, together with broad increases in glucuronidated and sulfated bile acid conjugates and depletion of unconjugated precursors. A similar pattern of elevated taurocholic and taurochenodeoxycholic acid in cord blood has been associated with small for gestational age (SGA) status^50^, and gastroschisis is itself associated with up to a 61% higher likelihood of SGA birth^51^. The bile acid signature observed in our cohort may therefore reflect the growth disturbance characteristic of this population and may indicate compromised enterohepatic circulation in the setting of intestinal exposure to amniotic fluid.

Heme catabolism by heme oxygenase yields biliverdin, which is reduced to bilirubin by biliverdin reductase^32^. In our cohort, biliverdin and all bilirubin isomers were reduced, with a non-significant trend toward heme accumulation, consistent with attenuated heme oxygenase activity. Unconjugated bilirubin functions as an endogenous antioxidant within the gastrointestinal tract and contributes to the maintenance of microbial richness and intestinal barrier integrity^52,53^. Reduced bilirubin in neonates with gastroschisis may therefore compromise antioxidant defense and intestinal homeostasis, processes already perturbed by the severe inflammation^54^ and reduced microbial diversity^55^ characteristic of this condition. Whether the heme and bilirubin signature represents a gastroschisis specific perturbation of heme oxygenase activity or an effect shared with other pregnancies complicated by preterm delivery and growth restriction cannot be resolved from the present cohort.

Fibrinopeptide A (5-16), a thrombin generated fibrinogen cleavage product considered a surrogate indicator of coagulation activity^56^, was markedly reduced in the gastroschisis group (FC = 0.17, *p* = 5 × 10⁻⁴). This finding suggests altered coagulation dynamics in the setting of gastroschisis^56^. Whether this reduction reflects diminished thrombin activity, altered hepatic fibrinogen synthesis, or accelerated peptide clearance in gastroschisis remains unclear and warrants further targeted investigation.

Several limitations should be acknowledged. The cohort is small (*n* = 23 per group), which limits statistical power to detect more subtle metabolic differences. The cases and controls also differed in gestational age at delivery and in the frequency of small for gestational age status, features that are themselves consequences of gastroschisis rather than independent characteristics of the infants; when the comparison was restricted to term births most of the signature was retained (Figure 7), although a residual gestational age difference persisted even within the term subset. The 53-metabolite signature is therefore reported as a cohort level finding that distinguishes gastroschisis from matched controls, and disentangling the effects specific to gastroschisis from those attributable to prematurity or growth restriction will require a larger cohort recruited with gestational age and birthweight matching. Additional limitations include the inability to resolve the maternal versus fetal origin of many detected metabolites from cord blood sampling alone, the cross sectional design at delivery which does not permit causal inference at the metabolite to outcome level, and the lipid bias of untargeted metabolomic reference libraries which may have biased identification away from amino acid and peptide pathways. Strengths of the study include prospective collection of cord blood, an unbiased untargeted profiling approach, and the largest cohort of this condition to date.

In summary, neonates with gastroschisis exhibit substantial alterations in cord blood lipid metabolism, steroidogenesis, bile acid conjugation, and heme catabolism. These signatures are concordant with the clinical phenotype of growth restriction, neurodevelopmental risk, and intestinal injury that characterizes this population. Direct comparison with published cord blood metabolomic signatures of small for gestational age, dominated by alterations in amino acid metabolism, lipoprotein subfractions, and acylcarnitines^57,58^, shows minimal overlap with our canonical signature, supporting the interpretation that the heme, ceramide, steroid hormone, and bile acid alterations identified here reflect gastroschisis biology rather than a generic growth restricted phenotype. A critical challenge will be to determine whether the metabolic perturbations identified here are causally linked to disease severity and long term outcomes, to define the molecular mechanisms by which gastroschisis remodels the fetal metabolome, and to confirm in a future cohort with gestational age and birthweight matched recruitment which of these signatures are gastroschisis specific and which are mediated through the prematurity and growth restriction that accompany the condition.

## Methods

### Study Participants

Participants were prospectively recruited from prenatal clinics and Labor and Delivery at Loma Linda University Children’s Hospital between 2015 and 2018. Cases of gastroschisis were identified by routine prenatal ultrasonographic examination and confirmed postnatally. Healthy controls were defined as singleton pregnancies without hypertensive disorders of pregnancy, diabetes mellitus, fetal congenital anomaly, or fetal growth restriction. Written informed consent was obtained from each participant; the study was approved by the Institutional Review Board at Loma Linda University (IRB# 5140043). Cases and controls were not matched a priori on maternal age or gestational age. The resulting imbalances and their effect on the differential abundance signal are reported in the demographic and sensitivity analyses below.

### Cord Blood Collection and Processing

At the time of delivery, cord blood was collected into EDTA containing tubes and transported directly to the clinical research laboratory at Loma Linda University Children’s Hospital by dedicated laboratory personnel maintained on call for this purpose. Plasma was isolated by centrifugation immediately upon arrival and stored at −80 °C until analysis.

### Untargeted Metabolomic Profiling

Cord blood plasma samples (*n* = 46) were submitted to Metabolon, Inc. (Morrisville, NC) for untargeted metabolomic profiling. Sample extraction was performed using a MicroLab STAR® automated liquid handling platform (Hamilton Company) and a 2000 Geno/Grinder (Glen Mills, Inc.). Multiple cocktail standard mixtures and extracted water samples were included throughout the workflow to ensure quality control. Sample extracts were divided into five fractions: two aliquots analyzed by reverse phase ultra performance liquid chromatography tandem mass spectrometry (UPLC-MS/MS) with positive ion electrospray ionization, one aliquot analyzed by reverse phase UPLC-MS/MS with negative ion electrospray ionization, one aliquot analyzed by hydrophilic interaction chromatography UPLC-MS/MS with negative ion electrospray ionization, and one aliquot retained as a backup sample. Residual organic solvents were removed by evaporation using a TurboVap® system (Zymark) followed by overnight storage under nitrogen. UPLC-MS/MS was performed as previously described^59^, using an ACQUITY UPLC system (Waters) coupled to a Q Exactive high resolution accurate mass spectrometer (Thermo Scientific) equipped with a heated electrospray ionization source and an Orbitrap mass analyzer operated at a mass resolution of 35,000. Each acquisition consisted of a full MS scan followed by data dependent MS/MS fragmentation, covering a mass to charge range of 70 to 1000.

### Compound Identification, Quantification, and Normalization

Raw MS data were processed for peak identification, and detected features were matched against a reference library of approximately 5,400 purified standards and 7,000 entries corresponding to structurally unnamed biochemicals. Compound mapping incorporated the Kyoto Encyclopedia of Genes and Genomes^60^, the Human Metabolome Database^61^, and PubChem^62^. Compound assignments required retention index matching within a defined tolerance window, accurate mass matching within ±10 ppm, and concordance of MS/MS fragmentation patterns evaluated by forward and reverse spectral matching scores. Metabolite abundance was quantified as the area under the chromatographic peak. Areas were normalized for run-to-run instrumental variability and for total protein content. Values were log transformed and rescaled such that the median value across samples for each metabolite was set to 1. Missing values attributable to technical limitations were imputed with the minimum observed value. Metabolites detected in fewer than 70% of the 46 participants (i.e., detected in 32 or fewer samples) were excluded from downstream analyses. After exclusion of xenobiotics, 767 endogenous metabolites were retained.

### Statistical Analysis and Visualization

Differential abundance was assessed by Welch’s two sample t test. Statistical significance was defined as p < 0.05; the false discovery rate was estimated using the q value method^63^. Pathway activation was calculated as the proportion of statistically significant metabolites within each sub_pathway. Unsupervised principal component analysis was performed using the prcomp function in R, and group separation was tested by permutational multivariate analysis of variance (PERMANOVA) using the vegan package^64^. Hierarchical clustering was performed using the pheatmap package^65^. Volcano plots and pathway dot plots were generated using dplyr^66^, ggplot2^67^, and ggrepel^68^. Analyses were performed in R Studio v2024.12.1+563^69^, ArrayStudio (Qiagen), and JMP Statistical Discovery (JMP). Schematic illustrations of metabolic pathways were prepared using BioRender.

## Notes

### Competing Interest Statement

The authors have declared no competing interest.

### Funding

- Loma Linday University Health, Department of Obstetrics and Gynecology

