## Supplementary Info for "Untargeted cord blood metabolomics reveals altered lipid metabolism in neonates with gastroschisis"

### Supplementary Figure

#### Supplementary Figure 1.

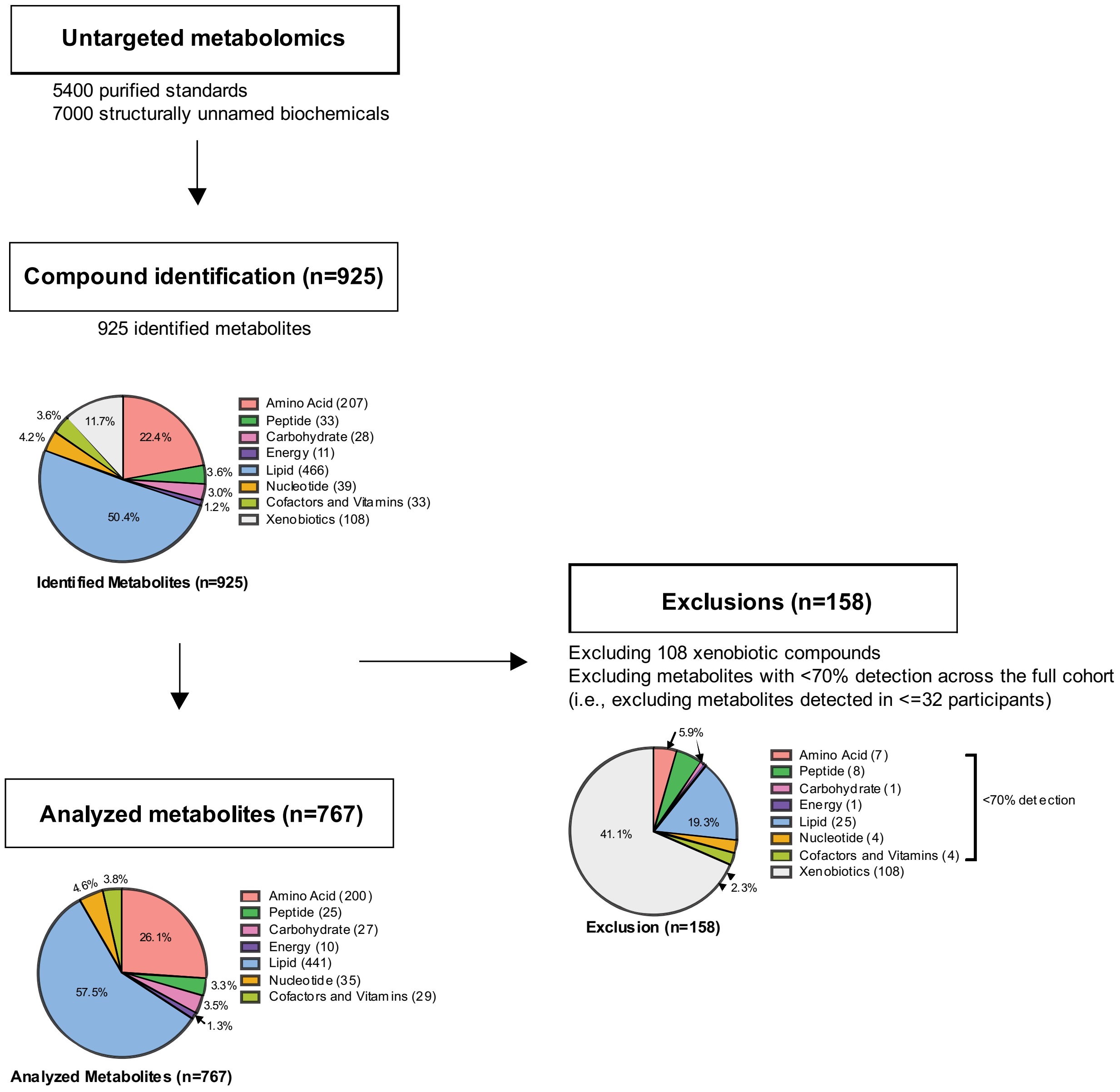


**Supplementary Figure 1.** Of approximately 5400 purified standards and 7000 structurally unnamed biochemicals in the Metabolon reference library, 925 metabolites were identified. After exclusion of all xenobiotics and metabolites detected in fewer than 70% of the cohort, 767 endogenous metabolites were retained for downstream analyses.

#### Supplementary Tables

**Supplementary Table 1.** Sensitivity of the 53 cohort significant metabolites to gestational age.

| Sub pathway | Biochemical | FC | p-value | q-value |
|---|---|---|---|---|
| **Hemoglobin and Porphyrin Metabolism** | biliverdin | 0.43 | $3.65 \times 10^{-5}$ | 0.0003 |
| | bilirubin (Z,Z) | 0.43 | 0.020 | 0.026 |
| | bilirubin (E,Z or Z,E) | 0.46 | 0.003 | 0.007 |
| | bilirubin (E,E) | 0.65 | 0.002 | 0.006 |
| **Polyamine Metabolism** | spermidine | 1.54 | 0.035 | 0.038 |
| | N-acetylputrescine | 1.45 | 0.0003 | 0.002 |
| | N1,N12-diacetylspermine | 1.43 | 0.002 | 0.005 |
| | 4-acetamidobutanoate | 1.32 | 0.002 | 0.005 |
| | 5-methylthioadenosine (MTA) | 1.32 | 0.034 | 0.037 |
| **Pyrimidine Metabolism** | orotate | 2.52 | $2.35 \times 10^{-5}$ | 0.0003 |
| | cytosine | 1.96 | 0.045 | 0.046 |
| | N-carbamoylaspartate | 1.49 | 0.004 | 0.009 |
| | 3-ureidopropionate | 1.46 | 0.007 | 0.012 |
| | 3-(3-amino-3-carboxypropyl)uridine | 1.35 | 0.0002 | 0.0009 |
| | N-acetyl-beta-alanine | 1.33 | 0.002 | 0.005 |
| | 2'-O-methylcytidine | 1.25 | 0.008 | 0.013 |
| | uridine | 1.24 | 0.007 | 0.011 |
| | orotidine | 1.23 | 0.035 | 0.038 |
| **Fibrinogen Cleavage Peptide** | fibrinopeptide A (5-16) | 0.17 | 0.0005 | 0.002 |
| | fibrinopeptide B (1-13) | 0.22 | 0.0002 | 0.001 |
| | fibrinopeptide A, des-ala(1) | 0.30 | 0.007 | 0.011 |
| | fibrinopeptide B (1-11) | 0.30 | 0.045 | 0.047 |

*For each metabolite the full cohort fold change and the fold change (FC) and p-value within term births are reported, together with whether the metabolite remained significant when the case versus control comparison was restricted to term births.*

**Supplementary Table 2. Lower fatty acid (FA) in neonates with gastroschisis.**

| Sub pathway | Biochemical | FC | p-value | q-value |
|---|---|---|---|---|
| Long Chain Saturated FA | myristate (14:0) | 0.59 | 0.0002 | 0.0013 |
| | pentadecanoate (15:0) | 0.69 | 0.0003 | 0.0016 |
| | palmitate (16:0) | 0.64 | 0.0025 | 0.006 |
| | margarate (17:0) | 0.62 | $6.14 \times 10^{-5}$ | 0.0005 |
| | stearate (18:0) | 0.70 | 0.0031 | 0.007 |
| | nonadecanoate (19:0) | 0.69 | $8.89 \times 10^{-5}$ | 0.0006 |
| | arachidate (20:0) | 0.82 | 0.0079 | 0.013 |
| Medium Chain FA | caproate (6:0) | 0.80 | 0.048 | 0.049 |
| | caprate (10:0) | 0.79 | 0.028 | 0.032 |
| | 10-undecenoate (11:1n1) | 0.76 | 0.019 | 0.025 |
| | laurate (12:0) | 0.57 | $7.10 \times 10^{-5}$ | 0.0005 |
| Long Chain MUFA | oleate/vaccenate (18:1) | 0.51 | 0.004 | 0.008 |
| | 10-nonadecenoate (19:1n9) | 0.56 | 0.0005 | 0.002 |
| | eicosenoate (20:1) | 0.57 | 0.017 | 0.024 |
| | 10-heptadecenoate (17:1n7) | 0.62 | 0.002 | 0.005 |
| Long Chain PUFA (n3 and n6) | linoleate (18:2n6) | 0.45 | 0.0004 | 0.002 |
| | linolenate (18:3n3 or 6) | 0.49 | 0.0004 | 0.002 |
| | eicosapentaenoate (EPA; 20:5n3) | 0.45 | 0.0003 | 0.002 |
| | docosapentaenoate (n3 DPA; 22:5n3) | 0.51 | $3.58 \times 10^{-5}$ | 0.0003 |
| | docosahexaenoate (DHA; 22:6n3) | 0.62 | 0.002 | 0.005 |
| | hexadecadienoate (16:2n6) | 0.61 | 0.001 | 0.004 |
| | dihomo-linoleate (20:2n6) | 0.63 | 0.0007 | 0.003 |
| | adrenate (22:4n6) | 0.63 | 0.021 | 0.027 |
| | nisinate (24:6n3) | 0.62 | 0.006 | 0.011 |
| | dihomo-linolenate (20:3n3 or n6) | 0.76 | 0.024 | 0.030 |
| | mead acid (20:3n9) | 1.32 | 0.014 | 0.020 |
| | docosatrienoate (22:3n6) | 1.99 | 0.001 | 0.004 |
| Branched Chain FA | (14 or 15)-methylpalmitate (a17:0 or i17:0) | 0.54 | $9.99 \times 10^{-6}$ | 0.0002 |
| | (16 or 17)-methylstearate (a19:0 or i19:0) | 0.50 | $7.22 \times 10^{-7}$ | $2.29 \times 10^{-5}$ |
| Fatty Acid, Dicarboxylate | hydroxy-CMPF | 0.21 | 0.005 | 0.010 |
| | octadecanedioate (C18-DC) | 0.54 | 0.002 | 0.005 |
| | eicosenedioate (C20:1-DC) | 0.56 | $3.63 \times 10^{-7}$ | $1.43 \times 10^{-5}$ |
| | azelate (C9-DC) | 0.57 | 0.002 | 0.005 |
| | 3-CMPFP | 0.58 | $1.31 \times 10^{-5}$ | 0.0002 |
| | glutarate (C5-DC) | 0.64 | 0.027 | 0.031 |
| | octadecenedioate (C18:1-DC) | 0.67 | 0.006 | 0.011 |
| | eicosanedioate (C20-DC) | 0.67 | 0.001 | 0.004 |
| | 4-hydroxy-2-oxoglutaric acid | 0.69 | 0.019 | 0.025 |
| | octadecadienedioate (C18:2-DC) | 0.73 | 0.014 | 0.020 |

| | | | | |
|---|---|---|---|---|
| | dodecanedioate (C12-DC) | 0.74 | 0.036 | 0.039 |
| | heptenedioate (C7:1-DC) | 0.75 | 0.029 | 0.033 |
| | 2-hydroxyadipate | 1.33 | 0.020 | 0.026 |
| **Fatty Acid, Monohydroxy** | 2-hydroxylaurate | 0.53 | $6.19 \times 10^{-8}$ | $5.68 \times 10^{-6}$ |
| | 13-HODE + 9-HODE | 0.61 | 0.0007 | 0.003 |
| | 3-hydroxystearate | 0.69 | 0.035 | 0.038 |
| | 3-hydroxypalmitate | 0.74 | 0.007 | 0.012 |
| | 2-hydroxymyristate | 0.74 | 0.003 | 0.007 |
| | 3-hydroxymyristate | 0.74 | 0.010 | 0.015 |
| | 2-hydroxystearate | 1.19 | 0.016 | 0.023 |
| | 2-hydroxyarachidate | 1.61 | $4.97 \times 10^{-5}$ | 0.0004 |
| | 2-hydroxybehenate | 1.60 | $7.95 \times 10^{-5}$ | 0.0006 |

*Fold change (FC) represents the ratio of mean metabolite levels in gastroschisis (n=23) relative to control (n=23). Statistical significance was assessed by Welch's two sample t test. Only metabolites with p < 0.05 are shown. Asterisks (*) denote metabolites identified with lower confidence.*

**Supplementary Table 3. Higher fatty acid downstream metabolism pathways in neonates born with gastroschisis.**

| Sub pathway | Biochemical | FC | p-value | q-value |
|---|---|---|---|---|
| Fatty Acid, Amide | margaramide (17:0) | 47.19 | 0.022 | 0.027 |
| | oleamide | 14.19 | 0.038 | 0.040 |
| | linolenamide (18:3) | 11.20 | 0.014 | 0.020 |
| | palmitamide (16:0) | 9.01 | 0.042 | 0.044 |
| | linoleamide (18:2n6) | 8.16 | 0.029 | 0.033 |
| | stearamide (18:0) | 6.63 | 0.015 | 0.021 |
| | eicosadienamide (20:2) | 4.41 | 0.006 | 0.011 |
| Acyl Carnitine, Long Chain Saturated | arachidoylcarnitine (C20) | 2.58 | $1.42 \times 10^{-8}$ | $1.73 \times 10^{-6}$ |
| | stearoylcarnitine (C18) | 1.80 | $4.79 \times 10^{-7}$ | $1.73 \times 10^{-5}$ |
| | lignoceroylcarnitine (C24) | 1.33 | 0.023 | 0.029 |
| | palmitoylcarnitine (C16) | 1.30 | 0.005 | 0.010 |
| | cerotoylcarnitine (C26) | 1.27 | 0.006 | 0.011 |
| | margaroylcarnitine (C17) | 1.22 | 0.031 | 0.035 |
| Acyl Carnitine, Monounsaturated | nervonoylcarnitine (C24:1) | 2.11 | 0.0001 | 0.0008 |
| | eicosenoylcarnitine (C20:1) | 1.89 | 0.002 | 0.006 |
| | ximenoylcarnitine (C26:1) | 1.84 | $8.26 \times 10^{-5}$ | 0.0006 |
| | cis-4-decenoylcarnitine (C10:1) | 1.42 | $5.97 \times 10^{-5}$ | 0.0005 |
| | oleoylcarnitine (C18:1) | 1.41 | 0.004 | 0.008 |
| | 3-decenoylcarnitine | 1.32 | 0.010 | 0.015 |
| | 5-dodecenoylcarnitine (C12:1) | 1.27 | 0.029 | 0.033 |
| Acyl Carnitine, Medium Chain | hexanoylcarnitine (C6) | 1.47 | 0.003 | 0.007 |
| | octanoylcarnitine (C8) | 1.37 | 0.001 | 0.004 |
| | decanoylcarnitine (C10) | 1.24 | 0.021 | 0.027 |
| Acyl Carnitine, Polyunsaturated | adrenoylcarnitine (C22:4) | 1.37 | 0.044 | 0.046 |
| | dihomo-linolenoylcarnitine (C20:3n3 or 6) | 1.36 | 0.010 | 0.015 |
| | dihomo-linoleoylcarnitine (C20:2) | 1.34 | 0.019 | 0.025 |
| | arachidonoylcarnitine (C20:4) | 1.27 | 0.026 | 0.031 |
| Acyl Choline | palmitoloelycholine | 2.39 | $9.91 \times 10^{-5}$ | 0.0007 |
| | dihomo-linolenoyl-choline | 1.96 | 0.0007 | 0.003 |
| | oleoylcholine | 1.69 | 0.0007 | 0.003 |
| | arachidonoylcholine | 1.50 | 0.004 | 0.008 |
| | palmitoylcholine | 1.46 | 0.004 | 0.008 |
| | stearoylcholine | 1.41 | 0.008 | 0.013 |
| Acyl Glycine | picolinoylglycine | 1.42 | 0.049 | 0.050 |
| | N-palmitoylglycine | 0.74 | 0.006 | 0.011 |

*Fold change (FC) represents the ratio of mean metabolite levels in gastroschisis (n=23) relative to control (n=23). Statistical significance was assessed by Welch's two sample t test. Only metabolites with $p < 0.05$ are shown.*

**Supplementary Table 4. Higher glycerophospholipid metabolism but lower plasmalogen in neonates born with gastroschisis**

| Sub pathway | Biochemical | FC | p-value | q-value |
|---|---|---|---|---|
| **Lysophospholipid** | 1-stearoyl-GPE (18:0) | 1.69 | $4.96 \times 10^{-6}$ | 0.0001 |
| | 1-palmitoyl-GPE (16:0) | 1.37 | 0.0011 | 0.0036 |
| | 1-oleoyl-GPE (18:1) | 1.58 | $1.48 \times 10^{-5}$ | 0.0002 |
| | 1-stearoyl-GPC (18:0) | 1.40 | 0.0024 | 0.006 |
| | 1-palmitoyl-GPC (16:0) | 1.26 | 0.003 | 0.0062 |
| **Phosphatidyl-choline (PC)** | 1-palmitoyl-2-oleoyl-GPC (16:0/18:1) | 1.26 | $3.20 \times 10^{-5}$ | 0.0003 |
| **Phosphatidyl-ethanolamine (PE)** | 1-palmitoyl-2-oleoyl-GPE (16:0/18:1) | 1.09 | 0.03 | 0.033 |
| **Plasmalogen** | 1-(1-enyl-palmitoyl)-2-linoleoyl-GPE (P-16:0/18:2) | 0.33 | $8.08 \times 10^{-5}$ | 0.0006 |
| **Plasmalogen** | 1-(1-enyl-stearoyl)-2-linoleoyl-GPE (P-18:0/18:2) | 0.29 | 0.0082 | 0.013 |

*Fold change (FC) represents the ratio of mean metabolite levels in gastroschisis (n=23) relative to control (n=23). Statistical significance was assessed by Welch's two sample t test. Representative significant metabolites are shown.*

**Supplementary Table 5. Higher ceramide metabolism in neonates born with gastroschisis.**

| Sub pathway | Biochemical | FC | p-value | q-value |
|---|---|---|---|---|
| **Ceramides** | N-stearoyl-sphingosine (d18:1/18:0) | 2.46 | $7.52 \times 10^{-7}$ | $2.29 \times 10^{-5}$ |
| | ceramide (d16:1/24:1, d18:1/22:1) | 2.14 | $8.75 \times 10^{-6}$ | 0.0002 |
| | ceramide (d18:1/20:0, d16:1/22:0, d20:1/18:0) | 1.84 | $3.29 \times 10^{-5}$ | 0.0003 |
| | N-stearoyl-sphingadienine (d18:2/18:0) | 1.77 | $3.66 \times 10^{-5}$ | 0.0003 |
| | N-palmitoyl-sphingosine (d18:1/16:0) | 1.72 | $8.73 \times 10^{-6}$ | 0.0002 |
| | ceramide (d18:1/17:0, d17:1/18:0) | 1.56 | 0.006 | 0.011 |
| | ceramide (d18:2/24:1, d18:1/24:2) | 1.53 | $2.33 \times 10^{-5}$ | 0.0003 |
| | N-palmitoyl-sphingadienine (d18:2/16:0) | 1.35 | 0.003 | 0.007 |
| **Dihydroceramides** | N-palmitoyl-sphinganine (d18:0/16:0) | 2.61 | $1.25 \times 10^{-7}$ | $7.62 \times 10^{-6}$ |
| | N-stearoyl-sphinganine (d18:0/18:0) | 2.50 | $3.29 \times 10^{-5}$ | 0.0003 |
| **Sphingomyelins** | sphingomyelin (d18:1/18:0, d18:0/18:1) | 1.29 | 0.015 | 0.021 |
| | sphingomyelin (d18:1/20:0, d16:1/22:0) | 1.22 | 0.047 | 0.048 |
| **Hexosylceramides (HCER)** | glycosyl-N-(2-hydroxynervonoyl)-sphingosine | 3.17 | 0.001 | 0.004 |
| | glycosyl ceramide (d18:1/20:0, d16:1/22:0) | 0.55 | 0.002 | 0.005 |
| | glycosyl-N-stearoyl-sphingosine (d18:1/18:0) | 0.66 | 0.006 | 0.011 |
| **Lactosylceramides (LCER)** | lactosyl-N-palmitoyl-sphingosine (d18:1/16:0) | 0.63 | 0.003 | 0.007 |
| | lactosyl-N-nervonoyl-sphingosine (d18:1/24:1) | 0.74 | 0.038 | 0.040 |

*Fold change (FC) represents the ratio of mean metabolite levels in gastroschisis (n=23) relative to control (n=23). Statistical significance was assessed by Welch's two sample t test. Only metabolites with p < 0.05 are shown.*

**Supplementary Table 6. Lower androgenic and estrogenic metabolism in neonates born with gastroschisis.**

| Sub pathway | Biochemical | FC | p-value | q-value |
|---|---|---|---|---|
| **Androgenic Steroids** | androsterone sulfate | 0.13 | 0.027 | 0.031 |
| | androstenediol (3β,17β) monosulfate (2) | 0.22 | $2.26 \times 10^{-6}$ | $5.51 \times 10^{-5}$ |
| | 5α-androstan-3β,17α-diol disulfate | 0.30 | $1.14 \times 10^{-5}$ | 0.0002 |
| | 5α-androstan-3β,17β-diol disulfate | 0.40 | $1.12 \times 10^{-5}$ | 0.0002 |
| | androstenediol (3β,17β) monosulfate (1) | 0.41 | 0.0003 | 0.001 |
| | androstenediol (3β,17β) disulfate (1) | 0.47 | 0.010 | 0.015 |
| | androstenediol (3α,17α) monosulfate (2) | 0.51 | 0.008 | 0.013 |
| | androsterone glucuronide | 0.55 | 0.004 | 0.008 |
| | 16α-hydroxy DHEA 3-sulfate | 0.56 | 0.016 | 0.022 |
| | androstenediol (3β,17β) disulfate (2) | 0.56 | 0.005 | 0.009 |
| | dehydroepiandrosterone sulfate (DHEA-S) | 0.60 | 0.002 | 0.005 |
| **Estrogenic Steroids** | estriol-3-glucuronide | 0.21 | $1.50 \times 10^{-7}$ | $7.84 \times 10^{-6}$ |
| | estrone 3-sulfate | 0.39 | 0.018 | 0.024 |
| | estriol 3-sulfate | 0.58 | 0.019 | 0.025 |

*Fold change (FC) represents the ratio of mean metabolite levels in gastroschisis (n=23) relative to control (n=23). Statistical significance was assessed by Welch's two sample t test. Only metabolites with $p < 0.05$ are shown.*

**Supplementary Table 7. Increased secondary bile acid metabolism in neonates born with gastroschisis.**

| Subpathway | Biochemical | FC | p-value | q-value |
|---|---|---|---|---|
| **Primary Bile Acid Metabolism** | chenodeoxycholate | 0.28 | $3.14 \times 10^{-10}$ | $1.15 \times 10^{-7}$ |
| | cholate | 0.56 | 0.028 | 0.032 |
| | taurocholate | 1.77 | 0.032 | 0.035 |
| **Secondary Bile Acid Metabolism** | deoxycholate | 0.34 | $6.52 \times 10^{-10}$ | $1.19 \times 10^{-7}$ |
| | lithocholate sulfate (1) | 0.44 | 0.016 | 0.022 |
| | hyocholate | 0.51 | 0.003 | 0.006 |
| | taurocholenate sulfate* | 2.26 | 0.0001 | 0.0008 |
| | taurochenodeoxycholic acid 3-sulfate | 2.59 | 0.005 | 0.010 |
| | taurolithocholate 3-sulfate | 3.25 | 0.026 | 0.030 |
| | glycodeoxycholate 3-sulfate | 3.54 | 0.014 | 0.020 |
| | tauroursodeoxycholic acid sulfate (1) | 4.22 | $1.47 \times 10^{-5}$ | 0.0002 |
| | taurodeoxycholic acid 3-sulfate | 4.38 | 0.0003 | 0.002 |
| | taurohyocholate* | 2.09 | 0.020 | 0.026 |

*Fold change (FC) represents the ratio of mean metabolite levels in gastroschisis (n=23) relative to control (n=23). Statistical significance was assessed by Welch's two sample t test. Only metabolites with $p < 0.05$ are shown.*
